# Integrated Model of the Vertebrate Augmin Complex

**DOI:** 10.1101/2022.09.26.509603

**Authors:** Sophie M Travis, Brian P Mahon, Wei Huang, Meisheng Ma, Michael J Rale, Jodi S Kraus, Derek J Taylor, Rui Zhang, Sabine Petry

## Abstract

Accurate segregation of chromosomes is required to maintain genome integrity during cell division. This feat is accomplished by the microtubule-based spindle. To build a spindle rapidly and with high fidelity, cells take advantage of branching microtubule nucleation, which exponentially amplifies microtubules during cell division. Branching microtubule nucleation relies on the hetero-octameric augmin complex, but understanding how augmin promotes branching has been hindered by a lack of structural information about the complex. Here, we report an integrated model of vertebrate augmin, combining cryo-electron microscopy, advanced protein structural prediction, and the visualization of fused bulky tags via negative stain electron microscopy. This strategy allowed us to identify the location and orientation of each subunit within the structure. Evolutionary analysis of augmin’s structure reveals that it is highly conserved across diverse eukaryotes, and that augmin contains a previously-unidentified microtubule binding site. Moreover, we identify homology with the kinetochore-localized NDC80 complex. This new model of the augmin complex provides insight towards the mechanism and evolution of branching microtubule nucleation.

## INTRODUCTION

Spindles are micron-scale microtubule (MT)-based structures that are built from scratch every time a eukaryotic cell divides. Spindle MTs are generated in a manner that is tightly spatially controlled, with their stable minus-ends pointing toward the spindle pole on the cell periphery and their growing plus-ends pointing toward the chromosomes at the spindle center. Critically, spindle MTs have to be continuously generated, as any given MT turns over within seconds, yet the spindle framework must often persist for up to an hour. Due to the spindle’s central importance for viability, multiple partially redundant pathways are used to generate spindle MTs. However, one pathway, branching MT nucleation, is responsible for generating the bulk of the MTs in the spindle^1,2,3^.

In branching MT nucleation, a new MT is nucleated from a side of an existing MT at a shallow angle^4,5,6^. Thus, MTs can be rapidly amplified while preserving their polarity. In the absence of branching, spindle formation is delayed, spindle MTs are missing in the spindle body, the spindle assembly checkpoint is activated, and cytokinesis cannot proceed^7,8,9,10^. Branching MT nucleation depends on recruitment of the MT template and nucleator, the γ-tubulin ring complex (γ-TuRC), to the side of a pre-existing mother MT to give rise to a branched MT with matching polarity to the mother ^6, 11,12,13^. Both recruitment of γ-TuRC, as well as maintenance of polarity^11,12,13,14^, are carried out by a conserved protein complex, augmin.

Augmin is an eight-subunit protein complex first discovered for its role in localizing γ-TuRC to cellular spindles. It was discovered independently in invertebrates^7^, vertebrates^8, 15^, and plants^16^. Vertebrate augmin consists of eight subunits, Haus1-8, that form two biochemically distinct subcomplexes, tetramers T-II and T-III. T-II binds MTs via the disordered N-terminus of Haus8^14, 17^, whereas T-III binds to γ-TuRC^14^ via the adaptor subunit NEDD1^13^, thus forming a bridge between the mother MT and the source of the branched MT. Negative stain electron microscopy established that the complex from both human and *X. laevis* forms an “h”-shape, with the bottom of the “h” sitting on the MT, and the stalk region pointing away from the mother MT^14, 17^. However, to date, no high resolution three-dimensional structural information has been available for either the complex as a whole or any fragments. In addition, lack of any identified structural homologs preclude hypotheses about augmin’s subunits and their organization within the complex, and, finally, it remains unclear how augmin interacts with either the MT or γ-TuRC to enable branching MT nucleation.

In this work, we report a medium resolution cryo-electron microscopy (cryo-EM) map of the *X. laevis* augmin complex. By leveraging recent advances in protein structural prediction, as implemented in AlphaFold-Multimer, and resolving structural ambiguities by tagging experiments coupled with negative stain electron microscopy, we report a molecular model of *X. laevis* augmin. Integrating this model with prior experiments, we identified the locations of the MT binding site, a putative secondary MT binding site, and the proposed γ-TuRC binding site, and demonstrated that the structure of augmin is highly conserved across eukaryotes. These results allow us to build, for the first time, a molecular model of the branch site, and, in addition, reveal insight into the shared evolutionary origin of augmin and the NDC80 kinetochore-localized complex.

## RESULTS

### Cryo-EM study of X. laevis augmin

The augmin complex is a flexible and extended assembly, and this dynamic nature makes augmin’s structure determination challenging. As the most suitable target for structural studies, we chose *X. laevis* augmin complex lacking the C-terminal domain of Haus6, which we previously reconstituted and purified^14^. Having recapitulated this purification strategy (Figure 1a), we next assessed augmin’s activity. We reconstituted branching MT nucleation in vitro by adding *X. laevis* augmin lacking the C-terminus of Haus6 and γ-TuRC purified using its activator, the γ-TuNA domain of CDK5RAP2^18^ to a GMPCPP-stabilized MT seed attached to glass. Upon addition of tubulin and GTP, new MTs nucleated from the side of the mother MT, implying that purified *X. laevis* augmin recruited γ-TuRC to the MT seeds and thus stimulated branching MT nucleation.. Purified *X. laevis* augmin is able to stimulate branching MT nucleation, thus, by implication, recruiting γ-TuRC to the MT seeds. Moreover, the branches formed at an acute angle from the MT seed, with nearly uniform polarity, and interestingly, were also able to recruit tubulin to the lattice. (Figure 1b, Figure S1). Thus, augmin lacking the C-terminus of Haus6 is biochemically active in its three main functions, namely binding MTs, recruiting γ-TuRC, and facilitating branching MT nucleation.

**Figure 1:**
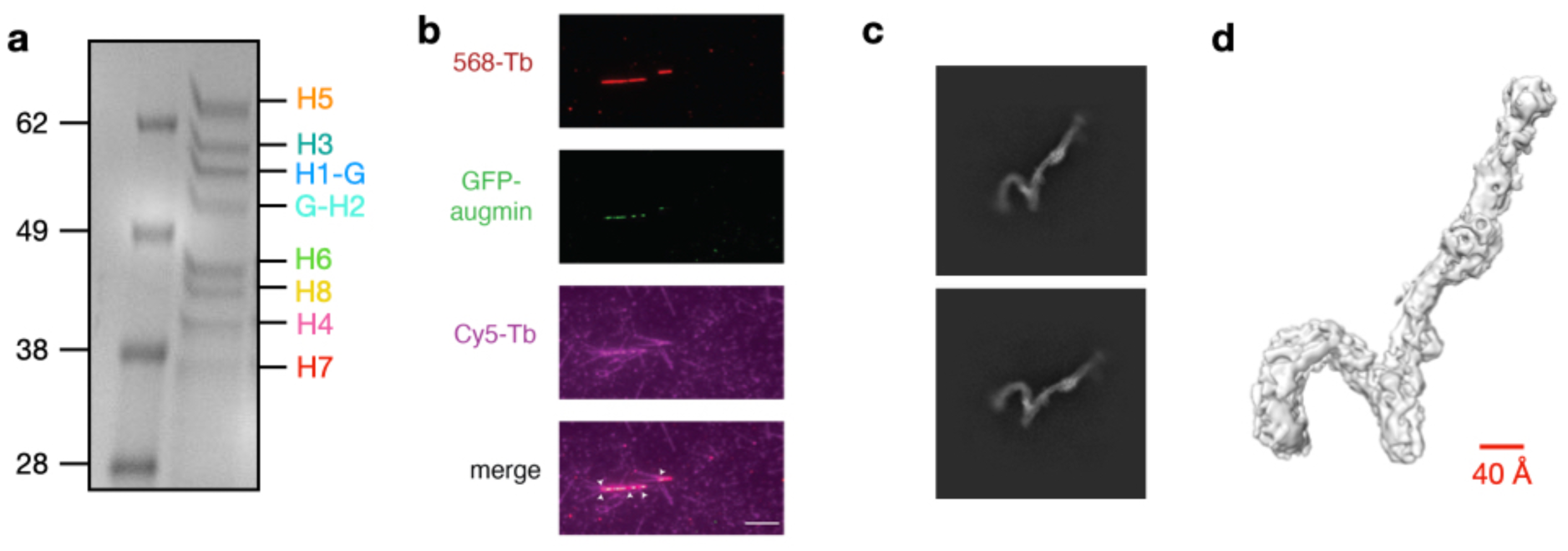
CryoEM structure of the *X. laevis* augmin complex. a. Stoichiometric *X. laevis* augmin octameric complex (lacking the C-terminus of Haus6) purified from insect cells. Here, and in subsequent figures, Haus1 through H8 is abbreviated as H1 through H8. b. Representative TIRF microscopy images from in vitro branching reconstitution with augmin and γ-TuRC. Alexa-568 labeled GMPCPP-stabilized MT seeds (red, top) were mixed with GFP-augmin and γ-TuRC allowing for localization of all components to the mother MT (green, 2nd panel) prior to initiation of in vitro branching MT nucleation. Branching MT nucleation was initiated by introducing 20 μM tubulin labeled with Cy5 (purple, bottom) whereby it was recruited to the GMPCPP-stabilized MT seeds (purple, bottom), and formed new MTs at an acute branch angle (arrowheads). Scale bars correspond to 5 μM. c. 2D class averages of the augmin complex in its preferred (front) orientation, demonstrating secondary structure. Box size is 650 Å. d. Post-processed electron density map of the augmin complex. Red line indicates a diameter measurement of 40 Å.

Next, we employed single-particle cryo-EM to unveil the structure for this challenging protein complex. We optimized the cryo-EM sample freezing condition, and found that Quantifoil holey carbon grids covered with additional continuous thin carbon film (∼5 nm thickness) produced the best mono-dispersed augmin particles. Using this strategy, we collected ∼3000 micrographs containing intact augmin particles. Due to the background noise of the added carbon, along with augmin being an inherently elongated particle, we encountered low particle contrast within our micrographs, which complicated data processing. Therefore, we used multiple strategies for particle picking, beginning with templates derived from negative stain data. Following 2D classification of the template picking results, we used the best particles to train a Topaz machine-learning algorithm^19^, which was then able to pick ∼100,000 intact augmin particles, resulting in 2D classes that displayed secondary structure features (Figure 1c). After 3D reconstruction and refinement, we obtained a cryo-EM reconstruction at 8.4 Å resolution (Figure 1d, Figure S2a, Table 1). The highest resolution region of the map was in the center of the T-III stalk (Figure S2b), and the map displayed slight anisotropy due to uneven angular distribution of the augmin particles on the EM grid (Figure S2c). At this resolution, we were able to establish that augmin has a relatively uniform radius across its entire length, comprising a cylindrical density of ∼40 Å in diameter (Figure 1d). This radius is consistent with that of a four-helix bundle, but, as 8.4 Å resolution is insufficient for de novo model building, we needed to incorporate structural information from orthogonal sources to further interpret the cryo-EM density.

**Table 1:** Cryo-EM data collection statistics.

|  | dataset1 | dataset2 |
| --- | --- | --- |
| Krios location | WUSTL | CWRU |
| Direct Detector | K2 | K3 |
| Cs (spherical aberration) (mm) | 0.01<sup>a</sup> | 2.7 |
| Magnification | 105,000 X | 64,000 X |
| Pixel size (Å) | 1.1 | 1.34 |
| Total electron dose (e<sup>-</sup>/Å<sup>2</sup>) | 84.3 | 66 |
| No. of images | 930 | 2,350 |

<sup>a</sup>The Titan Krios microscope at WUSTL is equipped with a Cs corrector.

### Integrated structural model of *X. laevis* augmin

The challenge of interpreting moderate-resolution density is an old one in structural biology, and much insight has been gained by docking high-resolution structural fragments or orthologous structures into low resolution maps (for example in the interpretation of the COPI vesicle coat^20^). However, this approach was not feasible for augmin because no high-resolution subcomplexes had previously been determined, nor had any structural homologues been identified. An unexpected solution presented itself last year with the groundbreaking progress in protein structural prediction heralded by the release of AlphaFold^21^ and, particularly, the updated algorithm optimized for multi-subunit complexes, AlphaFold-Multimer^22^ (as, for example recently implemented in interpretation of the nuclear pore complex^23^).

Although the first iteration of AlphaFold, which can model only individual subunits, predicted extended helical structures that were incompatible with the 8.4 Å map, we were able to gain much more insight from the updated AlphaFold-Multimer release. Leveraging our knowledge of the two tetrameric subcomplexes of augmin—T-II, comprising Haus2, and Haus6-8, and T-III, comprising Haus1, and Haus3-5—we were able to predict high-confidence structures of the two stable augmin tetramers that docked into the moderate resolution map (Figure 2a). AlphaFold-Multimer predicted that both tetramers fold into extended four-helix bundle structures, matching the radius observed in the map. Strikingly, it also predicted a set of insertions in the center of T-III matching the bulge observed in the center of the T-III stalk (Figure 2a). In addition to broadly agreeing with the 8.4 Å cryo-EM map, the AlphaFold-Multimer models recapitulated the predicted helical and entwined nature of subunits within the tetramers and were consistent with the assembly hierarchy previously determined by pull-downs and mass spectrometry^14^.

**Figure 2:**
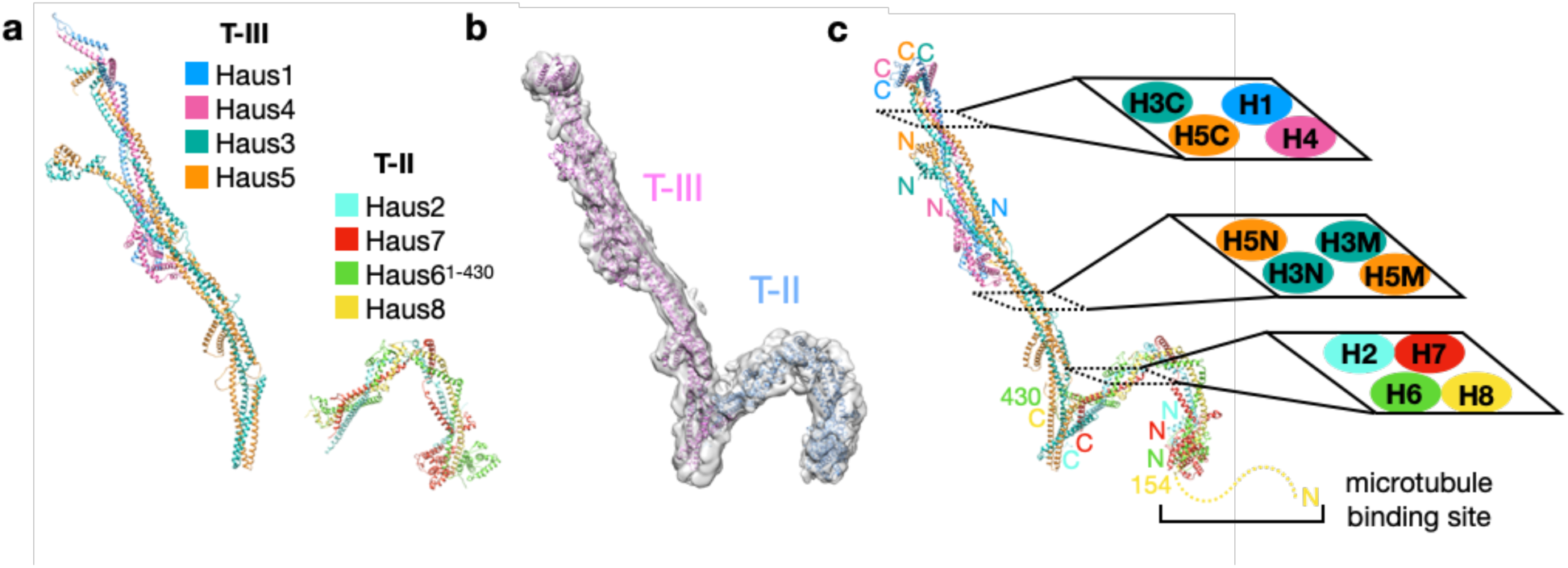
AlphaFold-Multimer based molecular model of the *X. laevis* augmin complex. a. Results of AlphaFold-Multimer prediction of *X. laevis* augmin subcomplexes: T-II (comprised of Haus2, Haus6^1–430^, Haus7, and Haus8) and T-III (comprised of Haus1, Haus3, Haus4, and Haus5). b. Final model of T-II and T-III into moderate resolution cryo-EM map shows good fit between predicted molecular contours and experimental data. c. Integrated model of the *X. laevis* augmin complex, highlighting predicted locations of N- and C- termini of all eight subunits, as well as the MT binding site present within the extreme disordered N-terminus of Haus8. Inset cross-sections through 4-helix bundles in T-II, T-III^core^, and T-III^ext^ show the architecture of the tetrameric parallel bundles within T-II and T-III^core^, as well as the antiparallel extended hairpin comprising T-III^ext^. Three helices derived from Haus3 appear in cross section, from the N-terminus (H3N) through the middle (H3M) to the C-terminus (H3C). Similarly, Haus5 appears in three cross-sections: N-terminal (H5N), middle (H5M), and C-terminal (H5C).

Two regions of the augmin model did not fit within the contours of our moderate resolution map and, thus, we employed molecular dynamics simulations with flexible fitting (MDFF) to better model these two sections (Figure S3). One section within T-III, formed by the extreme N-termini of Haus3 and Haus5, was predicted by AlphaFold-Multimer to form a second ‘leg’ emerging from the T-III central bulge. However, by negative stain, we see little evidence for this region in our 2D class averages (Figure 3a), and thus we expect this region to be mobile. In addition, the two globular domains at the extreme tip of T-II fit our map poorly, and MDFF predicts a combined rotation and translation of T-II at a hinge in the top of the arch, closing it by >40 Å and rotating the globular domains into density (Figure S3). As the density for this region is reasonably unambiguous, it seems likely that this mismatch is a result of imperfect prediction by AlphaFold-Multimer, although it could also herald dynamic motion in this region of augmin.

**Figure 3:**
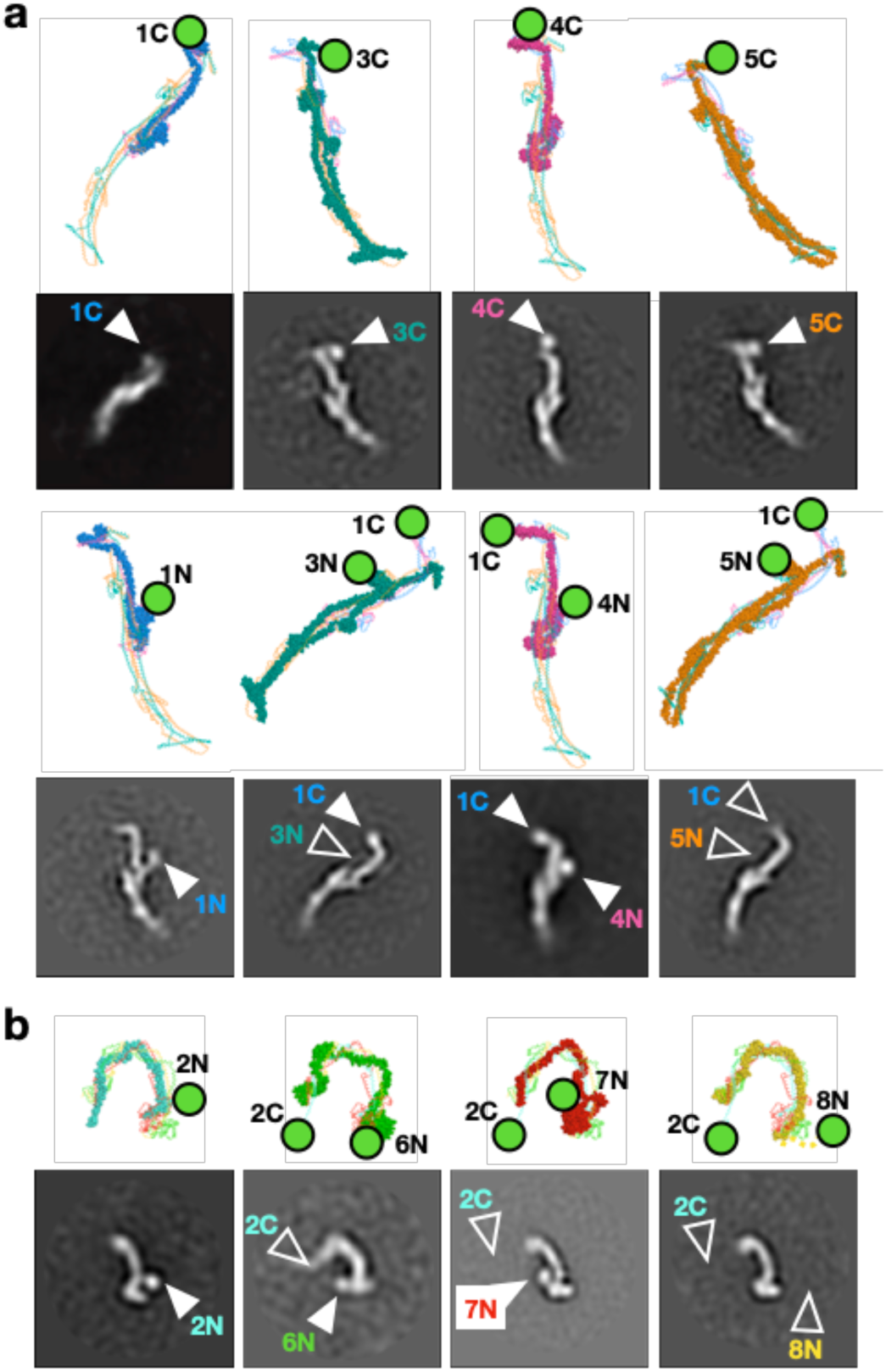
Negative Staining EM validation of augmin model. a. Eight T-III subcomplexes singly or doubly tagged with GFP. Above, predicted position(s) of GFP are annotated with green circles in the context of the T-III molecular model. Below, representative 2D negative stain classes are displayed for each complex. Visualized GFP density is indicated by solid arrows, whereas predicted but not visualized density is indicated by open arrows. Box size is 546 Å. b. Four T-II subcomplexes singly or doubly tagged with GFP. As above, predicted GFP position(s) are indicated in the top row, and experimentally observed 2D classes in the bottom. Visualized density is indicated with solid arrows, and predicted but not visualized density with open arrows.

The prediction of augmin’s structure by AlphaFold-Multimer provided insight into augmin’s global architecture. Perhaps unexpectedly, although T-II and T-III both consist of a core parallel 4-helix bundle and each is composed of a dimer of dimers, the global architecture of the two tetramers differs substantially. T-III^core^ is composed of paralogous subunits Haus1 and Haus4, which form a coiled-coil structure and intertwine with the coiled-coil C-termini of a second set of paralogs, Haus3 and Haus5 (Figure 2b, Figure S4a). The remaining three quarters of Haus3 and Haus5 form a continuous coiled-coil that folds back on itself into an extended antiparallel four-helix bundle, forming T-III^ext^, a bundle of equivalent length to T-III^core^ (Figure 2b). T-II is also comprised of two sets of paralogues, Haus6/7 and Haus2/8. However, in contrast with T-III^core^, in T-II paralogs do not form coiled coils with one another, but rather Haus6 and Haus8—structurally unrelated to one another—form a tight dimer that then associates with the Haus2/Haus7 dimer (Figure 2b, Figure S4b). Not only does T-III not share homology with T-II, but, following a DALI homology search^24^ of the AlphaFold predicted human proteome^25^, no obvious structural homologues could be found for any T-III subunits. However, the two T-II subunits Haus6 and Haus7 were unexpectedly predicted to contain globular calponin homology domains at their N-termini (Table 2), which will be discussed further below.

**Table 2:**
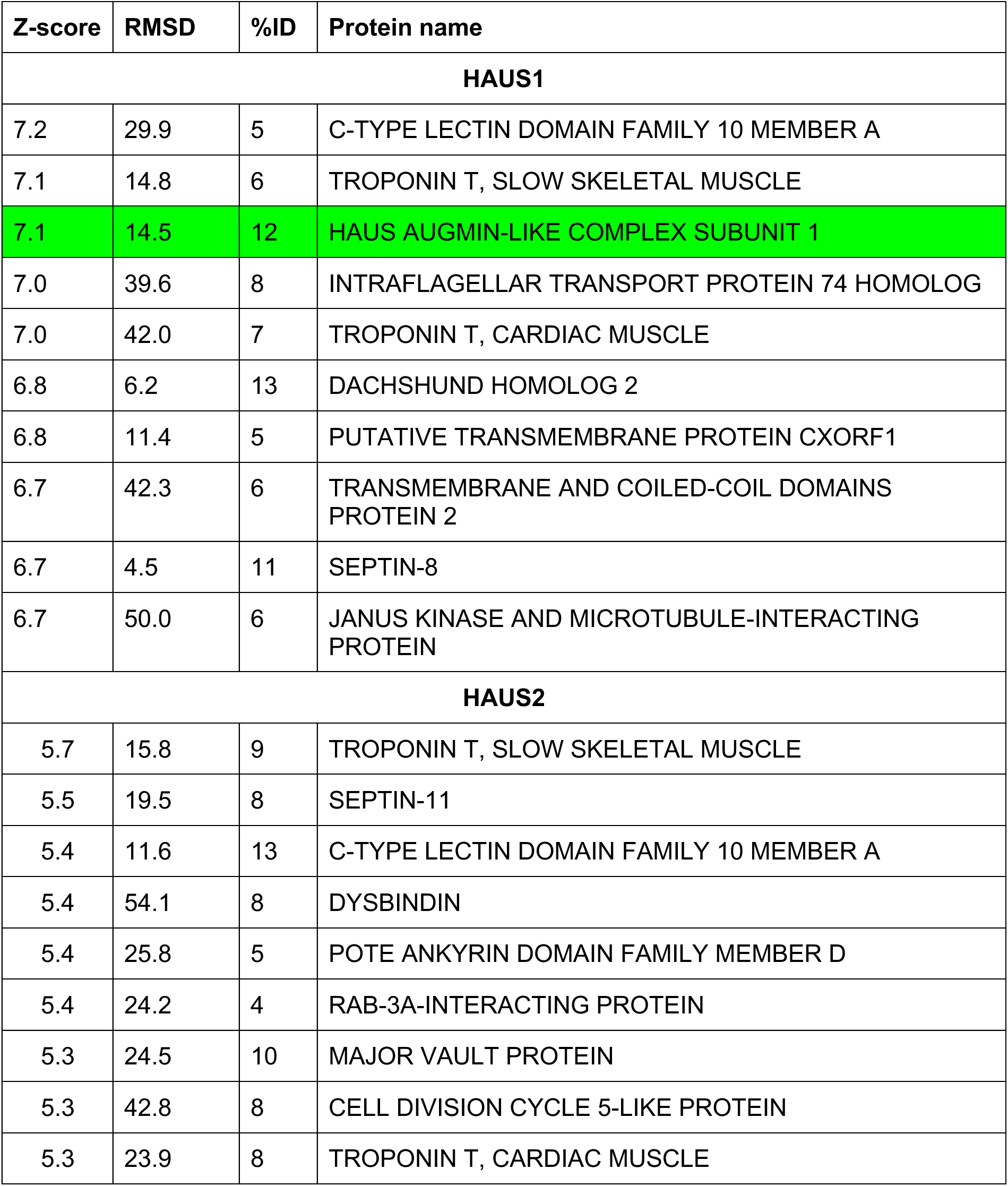
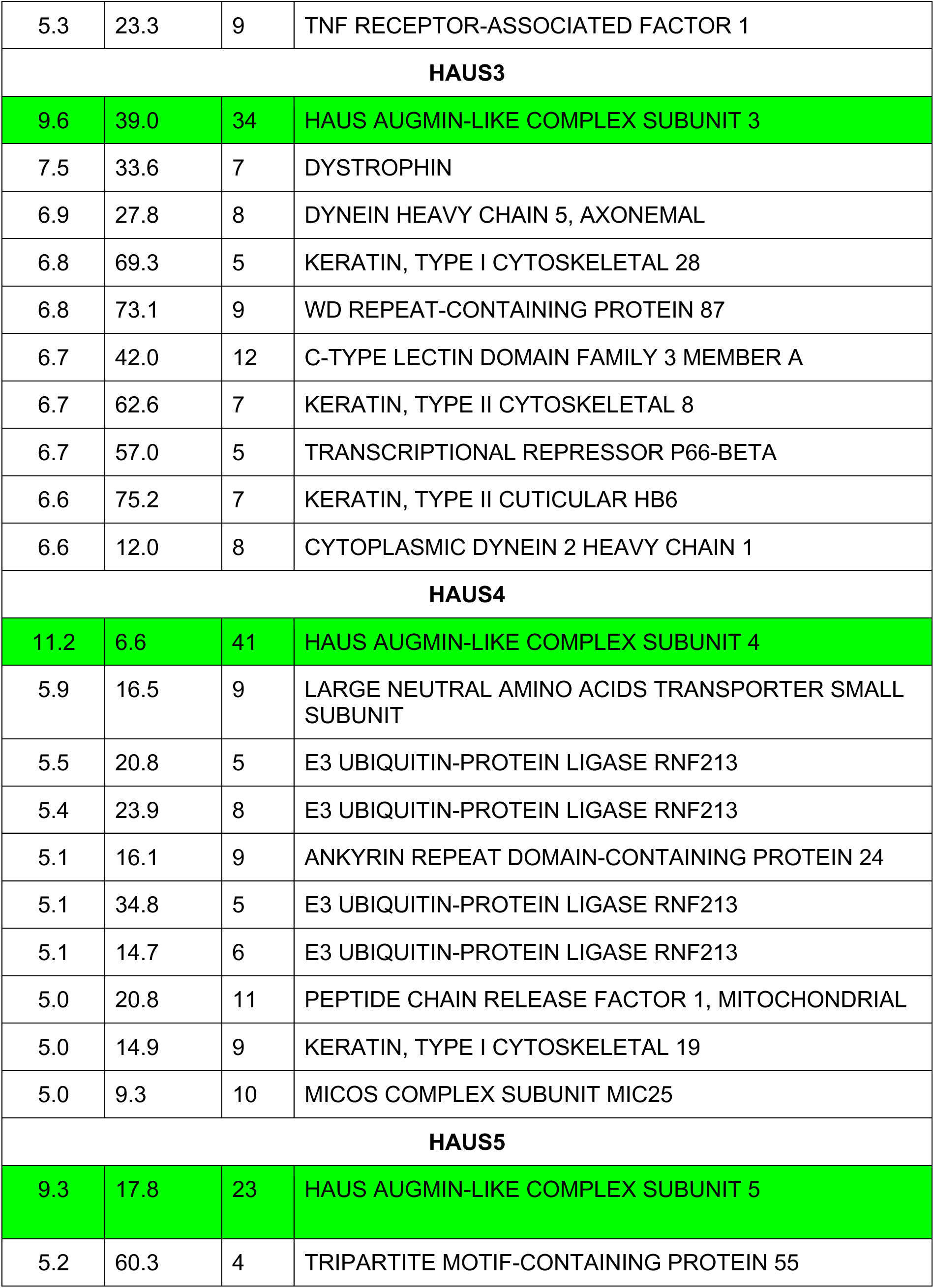
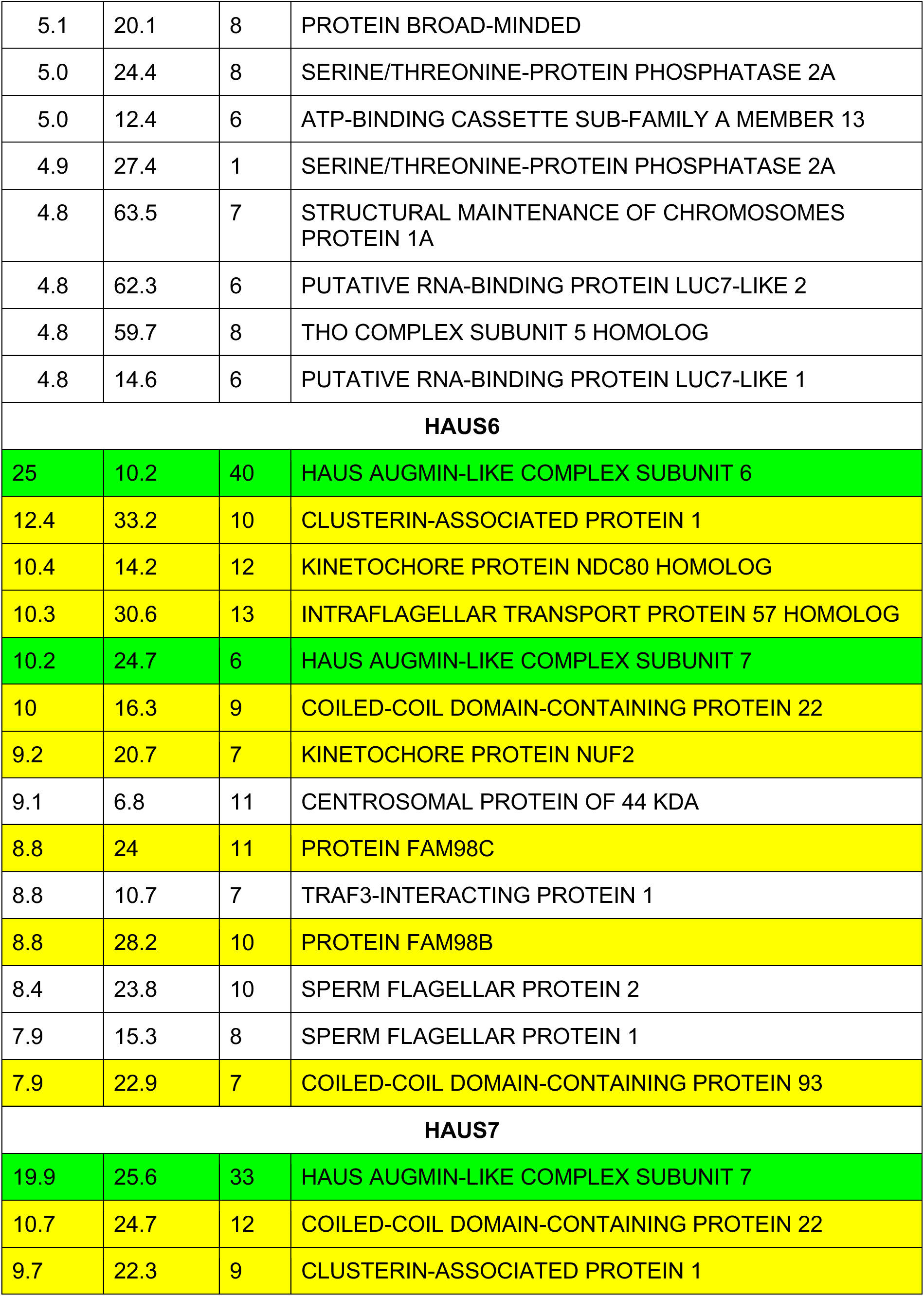
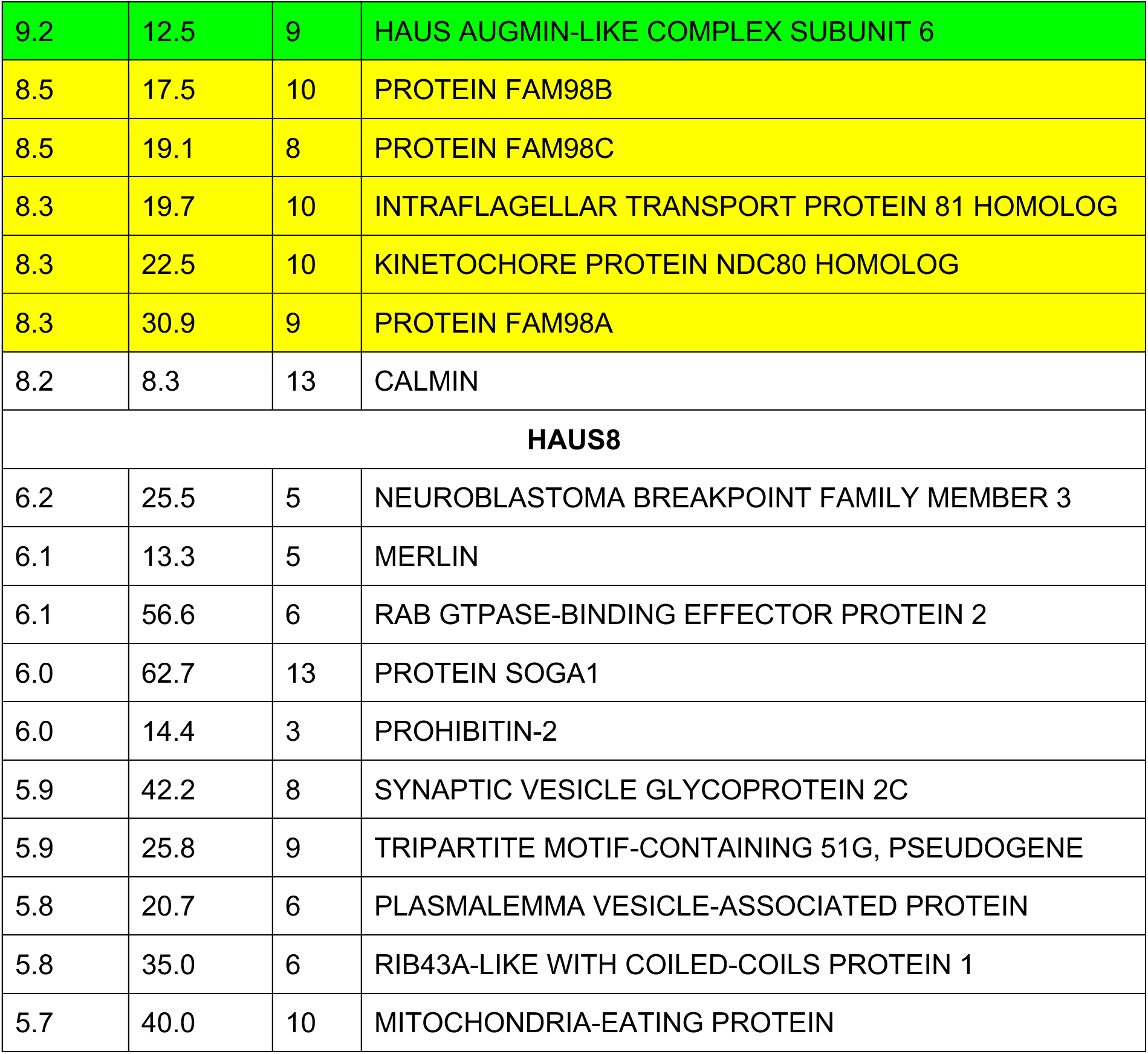
Top DALI-AlphaFold *H. sapiens* database hits for augmin subunits. The Z-score is DALI’s internal optimized similarity score, RMSD is the Cα root-mean-square displacement, and %ID is the pairwise sequence identity. Augmin subunits are highlighted in green, and NN-CH fold proteins in yellow

| Z-score | RMSD | %ID | Protein name |
| --- | --- | --- | --- |
| **HAUS1** |  |  |  |
| 7.2 | 29.9 | 5 | C-TYPE LECTIN DOMAIN FAMILY 10 MEMBER A |
| 7.1 | 14.8 | 6 | TROPONIN T, SLOW SKELETAL MUSCLE |
| 7.1 | 14.5 | 12 | HAUS AUGMIN-LIKE COMPLEX SUBUNIT 1 |
| 7.0 | 39.6 | 8 | INTRAFLAGELLAR TRANSPORT PROTEIN 74 HOMOLOG |
| 7.0 | 42.0 | 7 | TROPONIN T, CARDIAC MUSCLE |
| 6.8 | 6.2 | 13 | DACHSHUND HOMOLOG 2 |
| 6.8 | 11.4 | 5 | PUTATIVE TRANSMEMBRANE PROTEIN CXORF1 |
| 6.7 | 42.3 | 6 | TRANSMEMBRANE AND COILED-COIL DOMAINS PROTEIN 2 |
| 6.7 | 4.5 | 11 | SEPTIN-8 |
| 6.7 | 50.0 | 6 | JANUS KINASE AND MICROTUBULE-INTERACTING PROTEIN |
| **HAUS2** |  |  |  |
| 5.7 | 15.8 | 9 | TROPONIN T, SLOW SKELETAL MUSCLE |
| 5.5 | 19.5 | 8 | SEPTIN-11 |
| 5.4 | 11.6 | 13 | C-TYPE LECTIN DOMAIN FAMILY 10 MEMBER A |
| 5.4 | 54.1 | 8 | DYSBINDIN |
| 5.4 | 25.8 | 5 | POTE ANKYRIN DOMAIN FAMILY MEMBER D |
| 5.4 | 24.2 | 4 | RAB-3A-INTERACTING PROTEIN |
| 5.3 | 24.5 | 10 | MAJOR VAULT PROTEIN |
| 5.3 | 42.8 | 8 | CELL DIVISION CYCLE 5-LIKE PROTEIN |
| 5.3 | 23.9 | 8 | TROPONIN T, CARDIAC MUSCLE |
| 5.3 | 23.3 | 9 | TNF RECEPTOR-ASSOCIATED FACTOR 1 |
| **HAUS3** |  |  |  |
| 9.6 | 39.0 | 34 | HAUS AUGMIN-LIKE COMPLEX SUBUNIT 3 |
| 7.5 | 33.6 | 7 | DYSTROPHIN |
| 6.9 | 27.8 | 8 | DYNEIN HEAVY CHAIN 5, AXONEMAL |
| 6.8 | 69.3 | 5 | KERATIN, TYPE I CYTOSKELETAL 28 |
| 6.8 | 73.1 | 9 | WD REPEAT-CONTAINING PROTEIN 87 |
| 6.7 | 42.0 | 12 | C-TYPE LECTIN DOMAIN FAMILY 3 MEMBER A |
| 6.7 | 62.6 | 7 | KERATIN, TYPE II CYTOSKELETAL 8 |
| 6.7 | 57.0 | 5 | TRANSCRIPTIONAL REPRESSOR P66-BETA |
| 6.6 | 75.2 | 7 | KERATIN, TYPE II CUTICULAR HB6 |
| 6.6 | 12.0 | 8 | CYTOPLASMIC DYNEIN 2 HEAVY CHAIN 1 |
| **HAUS4** |  |  |  |
| 11.2 | 6.6 | 41 | HAUS AUGMIN-LIKE COMPLEX SUBUNIT 4 |
| 5.9 | 16.5 | 9 | LARGE NEUTRAL AMINO ACIDS TRANSPORTER SMALL SUBUNIT |
| 5.5 | 20.8 | 5 | E3 UBIQUITIN-PROTEIN LIGASE RNF213 |
| 5.4 | 23.9 | 8 | E3 UBIQUITIN-PROTEIN LIGASE RNF213 |
| 5.1 | 16.1 | 9 | ANKYRIN REPEAT DOMAIN-CONTAINING PROTEIN 24 |
| 5.1 | 34.8 | 5 | E3 UBIQUITIN-PROTEIN LIGASE RNF213 |
| 5.1 | 14.7 | 6 | E3 UBIQUITIN-PROTEIN LIGASE RNF213 |
| 5.0 | 20.8 | 11 | PEPTIDE CHAIN RELEASE FACTOR 1, MITOCHONDRIAL |
| 5.0 | 14.9 | 9 | KERATIN, TYPE I CYTOSKELETAL 19 |
| 5.0 | 9.3 | 10 | MICOS COMPLEX SUBUNIT MIC25 |
| **HAUS5** |  |  |  |
| 9.3 | 17.8 | 23 | HAUS AUGMIN-LIKE COMPLEX SUBUNIT 5 |
| 5.2 | 60.3 | 4 | TRIPARTITE MOTIF-CONTAINING PROTEIN 55 |
| 5.1 | 20.1 | 8 | PROTEIN BROAD-MINDED |
| 5.0 | 24.4 | 8 | SERINE/THREONINE-PROTEIN PHOSPHATASE 2A |
| 5.0 | 12.4 | 6 | ATP-BINDING CASSETTE SUB-FAMILY A MEMBER 13 |
| 4.9 | 27.4 | 1 | SERINE/THREONINE-PROTEIN PHOSPHATASE 2A |
| 4.8 | 63.5 | 7 | STRUCTURAL MAINTENANCE OF CHROMOSOMES<br>PROTEIN 1A |
| 4.8 | 62.3 | 6 | PUTATIVE RNA-BINDING PROTEIN LUC7-LIKE 2 |
| 4.8 | 59.7 | 8 | THO COMPLEX SUBUNIT 5 HOMOLOG |
| 4.8 | 14.6 | 6 | PUTATIVE RNA-BINDING PROTEIN LUC7-LIKE 1 |
| **HAUS6** |  |  |  |
| 25 | 10.2 | 40 | HAUS AUGMIN-LIKE COMPLEX SUBUNIT 6 |
| 12.4 | 33.2 | 10 | CLUSTERIN-ASSOCIATED PROTEIN 1 |
| 10.4 | 14.2 | 12 | KINETOCHORE PROTEIN NDC80 HOMOLOG |
| 10.3 | 30.6 | 13 | INTRAFLAGELLAR TRANSPORT PROTEIN 57 HOMOLOG |
| 10.2 | 24.7 | 6 | HAUS AUGMIN-LIKE COMPLEX SUBUNIT 7 |
| 10 | 16.3 | 9 | COILED-COIL DOMAIN-CONTAINING PROTEIN 22 |
| 9.2 | 20.7 | 7 | KINETOCHORE PROTEIN NUF2 |
| 9.1 | 6.8 | 11 | CENTROSOMAL PROTEIN OF 44 KDA |
| 8.8 | 24 | 11 | PROTEIN FAM98C |
| 8.8 | 10.7 | 7 | TRAF3-INTERACTING PROTEIN 1 |
| 8.8 | 28.2 | 10 | PROTEIN FAM98B |
| 8.4 | 23.8 | 10 | SPERM FLAGELLAR PROTEIN 2 |
| 7.9 | 15.3 | 8 | SPERM FLAGELLAR PROTEIN 1 |
| 7.9 | 22.9 | 7 | COILED-COIL DOMAIN-CONTAINING PROTEIN 93 |
| **HAUS7** |  |  |  |
| 19.9 | 25.6 | 33 | HAUS AUGMIN-LIKE COMPLEX SUBUNIT 7 |
| 10.7 | 24.7 | 12 | COILED-COIL DOMAIN-CONTAINING PROTEIN 22 |
| 9.7 | 22.3 | 9 | CLUSTERIN-ASSOCIATED PROTEIN 1 |
| 9.2 | 12.5 | 9 | HAUS AUGMIN-LIKE COMPLEX SUBUNIT 6 |
| 8.5 | 17.5 | 10 | PROTEIN FAM98B |
| 8.5 | 19.1 | 8 | PROTEIN FAM98C |
| 8.3 | 19.7 | 10 | INTRAFLAGELLAR TRANSPORT PROTEIN 81 HOMOLOG |
| 8.3 | 22.5 | 10 | KINETOCHORE PROTEIN NDC80 HOMOLOG |
| 8.3 | 30.9 | 9 | PROTEIN FAM98A |
| 8.2 | 8.3 | 13 | CALMIN |
| **HAUS8** |  |  |  |
| 6.2 | 25.5 | 5 | NEUROBLASTOMA BREAKPOINT FAMILY MEMBER 3 |
| 6.1 | 13.3 | 5 | MERLIN |
| 6.1 | 56.6 | 6 | RAB GTPASE-BINDING EFFECTOR PROTEIN 2 |
| 6.0 | 62.7 | 13 | PROTEIN SOGA1 |
| 6.0 | 14.4 | 3 | PROHIBITIN-2 |
| 5.9 | 42.2 | 8 | SYNAPTIC VESICLE GLYCOPROTEIN 2C |
| 5.9 | 25.8 | 9 | TRIPARTITE MOTIF-CONTAINING 51G, PSEUDOGENE |
| 5.8 | 20.7 | 6 | PLASMALEMMA VESICLE-ASSOCIATED PROTEIN |
| 5.8 | 35.0 | 6 | RIB43A-LIKE WITH COILED-COILS PROTEIN 1 |
| 5.7 | 40.0 | 10 | MITOCHONDRIA-EATING PROTEIN |

### Model validation through GFP-tagging and negative stain electron microscopy

Because structural prediction using AlphaFold is a comparatively recent development, we wanted to test the proposed augmin model using an established orthogonal technique that has previously been used to interpret the architecture of low-resolution electron microscopy structures^26, 27^. In order to independently establish the position and orientation of each subunit within T-III, we fused a bulky GFP tag, attached by a minimal linker, at the N- or C-terminus of each subunit of the tetramer. After purifying the tagged augmin tetramer complexes, we performed negative stain electron microscopy and looked for additional density corresponding to the GFP. Initially, C-terminal GFP tags were introduced to each of the four subunits, and we confirmed that all four C-termini localize to the end of the stalk, adjacent to one another (Figure 3a, top). Next, N-terminal GFP tags were introduced to the subunits, including a C-terminal GFP tag on Haus1 to aid in purification. Consistent with the AlphaFold-Multimer model, the N-termini of Haus1 and Haus4 localized to the center of the stalk, and the N-termini of Haus3 and Haus5 were not visualized, either due to flexibility of the termini and/or concealment behind the body of the stalk (Figure 3a, bottom). Similar experiments confirmed the location of the N-termini of T-II subunits Haus2, Haus6, and Haus7 at the globular-domain containing end of the tetramer (Figure 3b), although, due to flexibility of the other base of the T-II arch, GFP density on the C-termini of the T-II subunits could not be visualized. In addition, GFP tags localized to the extreme N-terminus of Haus8, the intrinsically-disordered primary MT binding site^17, 28^, yielded no additional observable density. In sum, visualization of fused bulky tags via negative stain electron microscopy helped validate our augmin model.

### Structural conservation of augmin across diverse eukaryotes

As discussed above, the augmin complex was first discovered in invertebrates (*D. melanogaster*)^7^, and only afterwards identified in vertebrates (*H. sapiens*)^8^ and plants (*A. thaliana*)^16^. Surprisingly, although a 1:1 correspondence has been established between vertebrate and plant augmin subunits – where for example HAUS1 in vertebrates is equivalent to AUG1 in plants, HAUS2 to AUG2, and so on – only 4 out of the 8 insect augmin subunits could be matched with their much more closely related vertebrate counterparts. For insects, the large subunits Dgt3 and Dgt5 have clear homology to Haus3 and Haus5, respectively, and Dgt6 and Dgt4 to Haus6 and Haus8. However, it was unclear to which vertebrate subunit the remaining four small insect subunits corresponded to, or even which tetramer they belonged in. Using AlphaFold-Multimer to predict the structure of *D. melanogaster* augmin, we were able to resolve this conundrum. We ran combinatorial predictions of all 12 possible *D. melanogaster* tetramers, containing either Dgt3/5 or Dgt4/6 and each possible pair of unassigned small subunits (Figure 4a). Only two predictions yielded solutions with structural homology to vertebrate augmin and thus we were able to equate Dgt2 with Haus4 (T-III), Dgt8 with Haus1 (T-III), Msd1 with Haus2 (T-II), and Msd5 with Haus7 (T- II).

**Figure 4:**
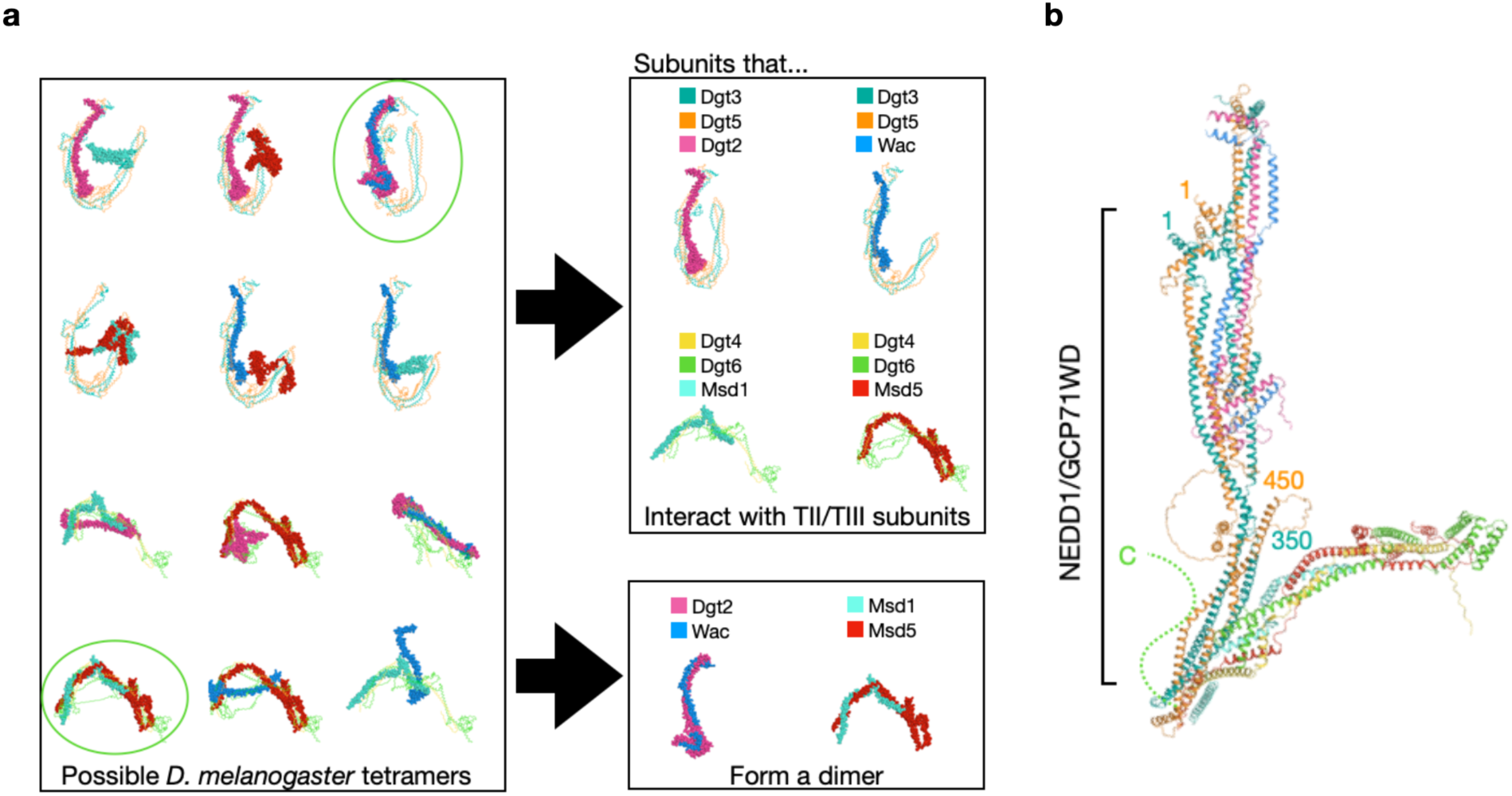
Homology-based Model of the *D. melanogaster* augmin complex. a. The structures of all six possible *D. melanogaster* T-III subcomplexes (top) and possible T-II subcomplexes (bottom) were predicted by AlphaFold-Multimer. Structures that superimpose with *X. laevis* T-II or T-III are circled. By categorizing the resulting predictions, Dgt2 and Wac were found to bind consistently with T-III subunits Dgt3 and Dgt5, as well as dimerizing with one another, and, similarly, Msd1 and Msd5 consistently dimerized with one another and bound T-II subunits Dgt4 and Dgt6. b. Structural alignment of the *D. melanogaster* subcomplexes T-II and T-III resulted in a molecular model of *D. melanogaster* augmin. The γ-TuRC adaptor protein GCP71WD (aka NEDD1)^29^, has been demonstrated to bind to the heterodimer of Dgt3^1–350^ and Dgt5^1–450^, and that span of residues, comprising the likely NEDD1 binding site, have been annotated.

Despite the low sequence similarity between *X. laevis* and *D. melanogaster* subunits, the predicted structure of *D. melanogaster* augmin well matched that of *X. laevis* augmin, particularly in the T-III subcomplex (Figure 4b). T-II displayed the same fold between both species, but *D. melanogaster* T-II was predicted to form a wider and shallower arch, due to a relative extension of its hinge. To further validate the *D. melanogaster* model, we integrated structural restraints previously obtained by the Wakefield group through cross-linking mass spectrometry (XLMS)^29^.

By incorporating these data into our structural model, we were able to demonstrate that AlphaFold-Multimer’s prediction of *D. melanogaster* augmin matched the majority of XLMS restraints, with the major exception of the only high-confidence cross-tetramer crosslink, between Msd1 Lys-113 and Dgt3 Ser-165 (Figure S5, Table 3). However, as the C-termini of all four *D. melanogaster* T-II subunits are also substantially shorter than their vertebrate counterparts, it seems likely that the interface between the two tetramers has coevolved to diverge from that found in vertebrate augmin.

**Table 3:** High-confidence *D. melanogaster* crosslinks and their predicted distances. BS3-crosslinks between the indicated residues are listed, along with their predicted distance in the AlphaFold-Multimer *D. melanogaster* augmin model (Figure S5). Crosslinks are derived from^29^. Disallowed cross-links (greater than 24 Å) are highlighted in yellow.

| Subunit 1 | Residue 1 | Subunit 2 | Residue 2 | Distance |
| --- | --- | --- | --- | --- |
| Dgt2 | Lys-193 | Dgt3 | Lys-508 | 18 Å |
| Dgt2 | Lys-55 | Dgt5 | Lys-154 | 12 Å |
| Dgt2 | Lys-214 | Dgt5 | Lys-653 | 9 Å |
| Dgt2 | Lys-193 | Wac | Lys-132 | 18 Å |
| Dgt2 | Lys-214 | Wac | Lys-146 | 21 Å |
| Dgt2 | Lys-214 | Wac | Lys-143 | 17 Å |
| Dgt2 | Lys-193 | Wac | Lys-140 | 19 Å |
| Dgt2 | Lys-217 | Wac | Lys-143 | 16 Å |
| Dgt2 | Lys-193 | Wac | Lys-143 | 27 Å |
| Dgt2 | Lys-217 | Wac | Lys-146 | 13 Å |
| Dgt3 | Lys-72 | Dgt5 | Lys-73 | 16 Å |
| Dgt3 | Lys-72 | Dgt5 | Lys-71 | 6 Å |
| Dgt3 | Lys-119 | Dgt5 | Lys-124 | 6 Å |
| Dgt3 | Lys-119 | Dgt5 | Lys-116 | 8 Å |
| Dgt3 | Lys-249 | Dgt5 | Lys-286 | 7 Å |
| Dgt3 | Lys-324 | Dgt5 | Lys-383 | 12 Å |
| Dgt3 | Lys-322 | Dgt5 | Lys-378 | 4 Å |
| Dgt3 | Ser-165 | Msd1 | Lys-113 | 120 Å |
| Dgt3 | Lys-508 | Wac | Lys-132 | 28 Å |
| Dgt4 | Lys-97 | Msd5 | Lys-174 | 20 Å |
| Dgt5 | Lys-625 | Wac | Lys-132 | 10 Å |
| Dgt5 | Ser-606 | Wac | Lys-132 | 30 Å |
| Dgt6 | Lys-270 | Msd1 | Lys-78 | 8 Å |
| Dgt6 | Lys-71 | Msd5 | Lys-87 | 21 Å |
| Dgt6 | Lys-143 | Msd5 | Lys-87 | 15 Å |
| Dgt6 | Lys-71 | Msd5 | Lys-86 | 23 Å |

In addition, with the *D. melanogaster* augmin model in hand, we were able to integrate further biochemical data about the location of binding sites on augmin for the γ-TuRC nucleator. Previous experiments have established the binding site of the γ-TuRC adaptor NEDD1—GCP71WD in *D. melanogaster*^29^. Chen and colleagues not only purified a heterodimer consisting of Dgt3^1–350^ and Dgt5^1–450^—roughly equivalent to T-III^ext^—they also demonstrated that this dimer was sufficient to bind to GCP71WD. Thus, we can put these facts together to map the NEDD1 binding location on *D. melanogaster* augmin to T-III^ext^ (Figure 4b). Intriguingly, this putative γ-TuRC binding location is adjacent to the unstructured C-terminus of Haus6, which, in vertebrates, has been proposed as the binding site of NEDD1^15, 30^. Therefore, regardless of the organism, γ-TuRC is likely to bind to the base of the T-III stalk.

Expanding upon our results with *D. melanogaster* augmin, we next sought to answer another major unknown question about the augmin complex, namely how well conserved it is across eukaryotes as a whole. Although no comprehensive analysis of augmin conservation is available, the complex has been reported missing from both budding and fission yeast, as well as nematodes^31^. To begin with, we performed a sequence-level search for homologs of the best-conserved subunit, Haus6, across the eukaryotic kingdom, and found that Haus6 was present in all five eukaryotic supergroups, although only well-conserved in metazoans and plants (Figure S6a). It remained unclear, however, whether the structure of the full complex was also conserved. Thus, we extended our prediction into representative members of each supergroup, with the exception of the SAR supergroup (*Straminopiles/Alveolata/Rhizaria)* where only Haus6 could be reliably identified in a single key species, *Tetrahymena thermophila*. Although, as with *Drosophila*, some small augmin subunits in these divergent eukaryotes could not be identified by sequence homology, structural prediction using all identified orthologues resulted in similar structures across *Opisthokonta*, *Plantae*, *Amoebozoa*, and *Discoba*, even among species with unidentified subunits (Figure S6b). This suggests that cryptic orthologs of the remaining small augmin subunits are likely present in these recalcitrant genomes and may perhaps be identifiable in the future using a 3D-structure-based search strategy such as DALI^24^ coupled with AlphaFold prediction of individual subunits. In addition, the augmin complex was apparently present in the last eukaryotic common ancestor and, although the full complex has been lost in many species since, where retained, augmin retains its structure, and thus perhaps its function.

## DISCUSSION

In this work, we have combined three orthogonal techniques—medium resolution cryo-EM, structural prediction using AlphaFold-Multimer, and subunit/subcomplex localization using negative stain electron microscopy—to assemble a molecular model of the *X. laevis* augmin complex. This model has allowed us not only to identify the positions and orientations of all eight subunits, but also to incorporate prior experimental knowledge to locate the MT and γ-TuRC binding sites within the complex, establishing for the first time the overall structure of the branching MT organizing center (Figure 5a). In addition, we have extended our modeling across diverse eukaryotic genomes and structurally identified augmin homologues, an identification that was not possible previously based on the sequence alone. We demonstrate that the predicted structure of the augmin complex is broadly conserved across four out of five eukaryotic supergroups, suggesting that augmin originated prior to the last eukaryotic common ancestor and that the complex’s function may remain broadly conserved.

**Figure 5:**
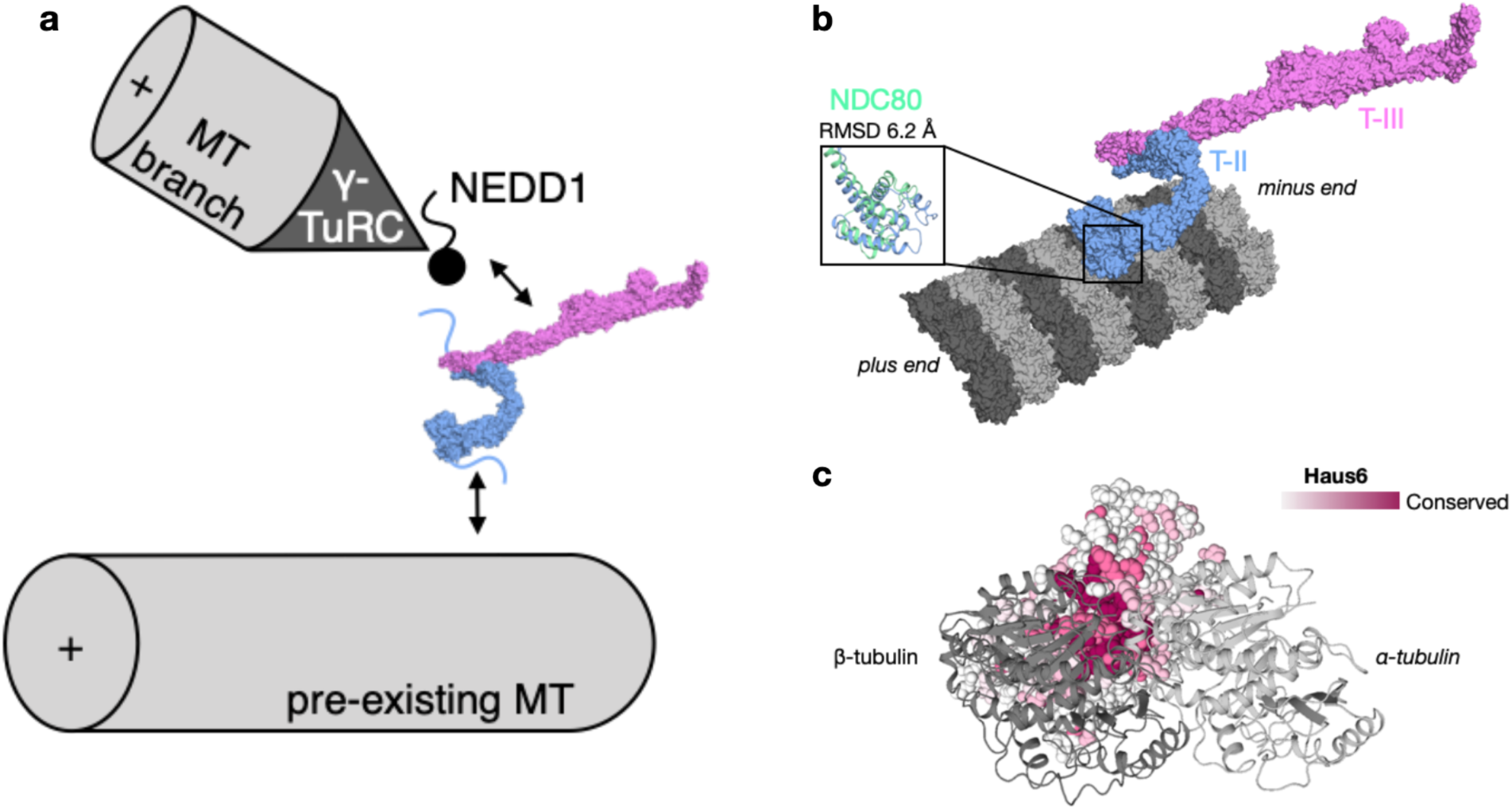
Interaction of augmin complex with the microtubule and other cellular factors. a. Overview of updated model of the MT branch site, including augmin molecular model. The NEDD1 γ-TuRC adaptor binding site is predicted to be located with T-III (pink), and MT binding site(s) within T-II (blue). b. Augmin T-II (blue)/T-III (pink) is superimposed onto the structure of the NDC80 complex (green) bound to the MT, with the superposition restricted to the calponin homology domains of Haus6 and Ndc80 (inset, with Cα RMSD displayed above). The NDC80 complex on the MT is derived from ^38^, PDB accession code 3IZ0. c. Augmin T-II, centered on the calponin homology of Haus6, is displayed as a surface colored by sequence conservation (where purple is the most conserved). A single tubulin dimer, positioned as in the NDC80-bound structure, is displayed in cartoon form (light gray, α-tubulin; dark gray, β-tubulin).

As we were preparing this manuscript, a model of *H. sapiens* augmin based on cryo-EM density and AlphaFold-Multimer structural predictions was published^32^. Our models are in general agreement, lending additional support to both. This is especially important since both models depend on the interpretation of moderate-resolution cryo-EM density using the cutting-edge but still relatively untested AlphaFold-Multimer modeling program. The largest difference between the two models lies in the relative hinging and rotation of the N-terminus of T-II, which, in the calponin homology domains, has moved ≥40 Å (Figure S4). This region was also the point of greatest difference between the *X. laevis* AlphaFold-Multimer model and the map, leaving open the question of whether the relative movement reflects intrinsic flexibility of this region of augmin or species-specific variability. In line with this second possibility, we note that the model of *D. melanogaster* augmin (Figure 4b) predicts a much wider T-II arch than either *H. sapiens* or *X. laevis* and that, as *D. melanogaster* MT branches extend at a wider angle than vertebrate branches^12, 33^, the relative hinging of T-II may be critical to establishing branching geometry.

In addition to shedding light on the conservation of augmin across eukaryotes, analysis of augmin’s predicted structure also has suggested intriguing new hypotheses about the complex’s origins and mechanism of action. As mentioned above, Haus6 and Haus7 displayed, unexpectedly, a classic calponin homology fold at their N-termini. This fold was first characterized as an actin binder, found in the calcium-dependent myosin regulator calponin. However, a divergent subfamily of calponin homology domains have shifted their affinity from actin to MTs, with the classic example being the MT plus-end binder EB1^34^. In addition to their similarity to all proteins containing tubulin-binding calponin homology domains, Haus6 and Haus7 displayed particularly close homology to members of the so-called NN-CH family (Table 2)^35^, a diverse group of MT binding multi-subunit complexes. This group includes the NDC80-complex subunits Ndc80 and Nuf2, which are homologs of Haus6 and Haus7. The NDC80 complex captures MTs near the kinetochore, which is also the area of highest augmin activity^36, 37^. Thus, it seems plausible that NDC80 and T-II might have diverged from a common ancestral complex deep in the evolutionary past.

In addition, by closely comparing the structures of NDC80 and augmin T-II, new insight can be gained into how augmin interacts with MTs. While the calponin-homology domain of Haus7 is poorly conserved on the sequence level and is missing altogether in some species including *D. melanogaster*, the calponin homology domain of Haus6 is one of the best-conserved regions of the augmin complex. Examining the predicted structure of Haus6’s calponin homology domain more closely, it becomes clear that one face of the domain presents a highly conserved basic surface (Figure 5b). Upon alignment with the calponin homology domain of the Ndc80 subunit, it is apparent that the conserved Haus6 surface overlaps with the MT binding face of Ndc80’s calponin homology domain, the primary MT binding site of the complex^38^. Although the primary MT binding site of augmin has been shown to reside within the disordered N-terminal 150 residues of Haus8^17^, previous results have suggested that T-II harbors a second MT binding site^17^ and, as the calponin homology domain of Haus6 is adjacent to the N-terminus of Haus8, it seems probable that these two regions could, together, form a bipartite MT binding site to stabilize T-II on the MT lattice. This second MT binding mode may help solve the mystery of how a disordered attachment via the unstructured N-terminus of Haus8 is capable of orienting γ-TuRC stably relative to the pre-existing MT, and more generally how branching MT nucleation is capable of maintaining spindle polarity.

## MATERIALS AND METHODS

### Protein expression & purification

Expression of the wild-type augmin Haus6^1–430^ complex has previously been described from Sf9 cells^14^. GFP-tagged versions of augmin subunits were generated by subcloning individual augmin subunits into custom pFASTBAC vectors containing either N-terminal TwinStrep-GFP or C-terminal GFP-TwinStrep tags. Bacmids were then generated by transformation into DH10Bac (NEB) *E. coli*, and subsequent screening on XGAL colorimetric LB plates. Bacmids were purified from DH10Bac culture, screened for insertion by polymerase chain reaction, and transfected into Sf9 cells using standard procedures. Tagged viral stocks were substituted for untagged stocks during co-infection of Sf9 cells with individual viruses bearing each of the eight subunits for augmin co-expression.

Augmin complexes and subcomplexes were purified as previously described^14^. 1 L infected Sf9 cells (for T-II and T-III subcomplexes) or 2 L cells (for octameric augmin) were harvested at 500x g and resuspended in lysis buffer (50 mM Tris pH 7.7, 200 mM NaCl, 5 mM EDTA, 3 mM β- mercaptoethanol, 10% glycerol, 0.05% Tween 20) supplemented with 10 ug/mL DNase I (Roche) and 1 cOmplete protease inhibitor cocktail tablet (Roche). After lysis via Emulsiflex, lysates were clarified at 200,000 g and loaded onto either 1 mL StrepTactin Superflow (IBA), for T-III, or 1 mL IgG Sepharose (Cytiva), for T-II and full augmin complex. After 3 hours batch binding, resin was washed with 100 column volumes of lysis buffer and eluted either with 2.5 mM desthiobiotin (for T-III) or by overnight cleavage with PreScission protease (for T-II and full complex). Eluates were concentrated using 50 kDa MWCO concentrators (Amicon) and further purified over a Superose 6 Increase 10/300 pre-equilibrated in CSF-XB (10 mM HEPES pH 7.7, 100 mM KCl, 5 mM EGTA, 2 mM MgCl2, 1 mM DTT). Peak fractions were analyzed for purity by SDS-PAGE, pooled, and flash frozen for -80°C storage. *X. laevis* γ-TuRC was isolated from *X. laevis* egg extract using a purification strategy available in preprint^18^. Here we utilized magnetic beads (cat. #G7287, Promega, Madison, WI) containing a pre-bound Halo-tagged CM1-containing peptide, Halo-γ-TuNA, to affinity purify native γ-TuRC from egg extract. γ-TuRC was eluted from magnetic beads via protease cleavage using a PreScission protease in a single fraction. The γ-TuRC elution was then concentrated using a 100 kDa MWCO Amicon 4 mL spin concentrator, and further purified with a 10-50% (w/w) sucrose gradient spun in a TLS55 rotor with a Beckman Coulter Optima MAX-XP ultracentrifuge at 200,000g, 2°C for 3h. The sucrose gradient was fractionated and each fraction was analyzed using western blot and negative-stain EM to determine the peak, which contained 100-150 nM purified γ-TuRC. Purified γ-TuRC was stored in CSF-XB buffer (100 mM KCl, 10 mM K-HEPES, 1 mM MgCl2, 0.1 mM CaCl2, 5 mM EGTA, pH 7.7) and ∼30% w/w sucrose.

### In vitro Branching MT Nucleation reaction using TIRF microscopy

In vitro reconstituted branching MT nucleation reactions were performed as previously described^11, 18^. In brief, we first functionalized coverslips with biotin-CONH-PEG-NH2 (cat. #103000-20 and #133000-25-20, Rapp Polymere, Tübingen, Germany), using the procedure described by Surrey and coworkers^39^. To make imaging chambers for TIRF microscopy assays, we used double-sided tape to outline channels on passivated glass slides, which we could then sandwich with functionalized coverslips.

For in vitro branching MT nucleation reactions, a mixture of augmin-HAUS6^1–430^ (50 nM) and γ- TuRC (10 nM), was first incubated for 5 min on ice, and then flowed into the imaging chamber which contained single cycled, biotin labeled Alexa-568 GMPCPP-stabilized MTs. Components were allowed to incubate within the imaging chamber for 5 min at room temperature to allow for localization of all branching components to stabilized MTs prior to initiating the reaction. After incubation, unbound proteins were removed by washing the reaction chamber with BRB80 (80 mM PIPES, 1 mM MgCl2, 1 mM EGTA, pH 6.8 with KOH). To initiate branching MT nucleation reactions, the final assay mixture was flowed into the chambers which consisted of the following components: 80 mM PIPES pH 6.8, 30 mM KCl, 1 mM EGTA, 1 mM MgCl2, 1 mM GTP, 5 mM β-mercaptoethanol, 0.075% (w/v) methylcellulose (4000 cP), 1% (w/v) glucose, 0.02% (v/v) Brij-35, 250 nM glucose oxidase, 64 nM catalase, 1 mg/ml bovine serum albumin, 19 μM unlabeled bovine tubulin and 1 μM Cy5-labeled bovine tubulin. Once assay mixture was added, the reaction was quickly transferred to the microscope stage and imaged using TIRF.

TIRF microscopy was performed with a Nikon TiE microscope using a 100 × 1.49 NA objective. Andor Zyla sCMOS camera was used for acquisition, with a field of view of 165.1 × 139.3 µm, multi-color images were acquired using NIS-Elements software (Nikon). All adjustable imaging parameters (exposure time, laser intensity, and TIRF angle) were kept the same within experiments. During data acquisition of in vitro branching MT nucleation reactions, the TIRF objective was warmed to 33°C using an objective heater (Bioptechs, 150819–13), and data was collected using time-lapse imaging, multi-color images collected every 2 sec. ImageJ software was used for image processing and data analysis.

### EM data collection

Negative-stained EM samples were prepared by diluting purified augmin to 150 nM in CSF-XB and pipetting 3 µl onto glow-discharged (15 mA, 25-30 secs) carbon film, 400 mesh Cu grids (Electron Microscopy Sciences), staining with 0.75% uranyl acetate solution. Negative-stain EM data was collected at 94,000x magnification (1.56 Å/pixel) with single-tilt using a Talos F200X Transmission Electron Microscope equipped with a 4k x 4k Ceta 16M CMOS camera.

Cryo-EM grids were prepared similarly using undiluted, purified augmin. 0.05% NP-40 was added to augmin prior to applying to grids. Here, 3 µl of sample was applied to glow-discharged (10 mA, 8 secs) Quantifoil holey carbon R 1.2/1.3 400 mesh grids coated with a home-made thin carbon film (∼5 nm thickness) using Leica EM ACE600 High Vacuum Sputter Coater. The grids were flash frozen in liquid ethane using a FEI Vitrobot Mark IV (Thermo Scientific) plunge freezer, using a blot force of 0 and with a 4.5 sec blot time. Cryo-EM data were collected using the Titan Krios microscopes at either Washington University in St. Louis (WUSTL) or Case Western Reserve University (CWRU). The data collection parameters are listed in Table 1.

### EM data processing

Data processing of negative-stain EM data was done using Relion^40^. Here, raw micrographs were used to manually pick particles for alignments and averaging. ∼10,000 total particles were manually picked for each tagged complex, followed by particle extraction and 3-10 rounds of 2D class averaging.

Data processing of cryo-EM data was done using CryoSparc (Structura Biotechnology). Raw micrograph movies were motion corrected and CTF-corrected using CryoSparc Live’s motion correction algorithm and PatchCTF, respectively. Augmin templates were generated from negative stain class averages of full length augmin and augmin T-III and independently used to pick particles, in order to account for model bias from template picking. After particle extraction and 2D classification, both sets of classes displayed the characteristic “h” shape of the full augmin complex. Particles from the best 3 classes (amounting to 15,000 particles) were then input to the Topaz machine learning particle picking algorithm^19^ using the ResNet8 architecture. Topaz was then used to pick ∼100,000 particles, which were sorted by 2D classification, and the best 30,000 were used for de novo 3D model generation. Next, the ∼60,000 good particles selected from 2D classification were used for two rounds of 3D refinement with dynamic masking. The resulting map was post-processed by the deep learning algorithm DeepEMhancer^41^, using the “tightT” setting. Local resolution was determined in CryoSparc.

### AlphaFold structural prediction and model docking

Canonical isoforms and sequences of *X. laevis* augmin subunits were input either singly into AlphaFold 2.1^21^ or in multimeric groups, using the --*multimer* option. The resulting five multiple models were compared for structural convergence and only converging models considered. Each of the five independent T-II and T-III models were rigid-body docked into the cryo-EM map and its inverse-hand equivalent using Chimera Fit Into Map. Then T-II was manually shifted away from T-III at their interfaces to avoid steric clashes. Input files for all-atom MDFF simulations were prepared starting from this structure with AutoPSF, mdff^42^ and restraints packages in VMD1.9.6^43^. MDFF simulations were performed with gscale = 0.3 and sampled for 3.825 ns using NAMD3 on a single RTX3060 GPU card. Cross-correlation values for snapshots every 1 ps from the MD trajectory with cryo-EM map were calculated to monitor the convergence of the fitting. The last frame of the total 3.825 ns simulation with a cc value of 0.8 was selected and subsequently refined in phenix^44^ with minimization_global option to reduce rotamer outliers. Structural alignments were performed using Cα superposition in PyMol. Surface conservation of T-II was calculated using the ConSurf server^45^. Structure figures were generated in either Chimera^46^ or PyMol (Schrodinger).

Orthologues of augmin subunits were identified through a combination of literature review, BLAST search using the domain enhanced lookup time accelerated-BLAST algorithm^47^, and HMMR hidden Markov model search^48^. Orthologues that had not been experimentally verified were validated for sequence completeness by alignment to their closest 10 homologs via BLAST search in the full UniProt database, and verified as non-spurious augmin orthologs by targeted BLAST search in the *X. laevis* or *A. thaliana* genome to ensure that the expected augmin subunit was the top hit. Structure-based homology search was performed using the DALI server^24^, searching either the full PDB experimentally determined database or the full *H. sapiens* AlphaFold predicted genome^25^.

## ACKNOWLEDGEMENTS

We would like to thank current and former members of the Petry lab for support and scientific feedback, particularly Collin McManus and Venecia Valdez for critical reading of the manuscript. We thank Michael Rau and James Fitzpatrick at WUCCI and Kunpeng Li at CWRU for microscopy support. We are grateful to Fred Hughson, Phil Jeffrey, and Alan Brown for helpful scientific discussions. We would also like to thank the staff of the Princeton Imaging and Analysis Center, particularly John Schreiber, for their technical assistance in electron microscopy, and Matthew Cahn of Princeton Research Computing for his assistance with data processing and AlphaFold implementation.

This work was supported by National Institutes of Health grants F32GM142149 (SMT), the Helen Hay Whitney Foundation (JK), NIGMS grant 1R01GM138854 (RZ), and NIGMS grant R01 1R01GM141100-01A1 (SP). The authors acknowledge the use of Princeton’s Imaging and Analysis Center (IAC), which is partially supported by the Princeton Center for Complex Materials (PCCM), a National Science Foundation (NSF) Materials Research Science and Engineering Center (MRSEC; DMR-2011750). Molecular graphics and analyses performed with UCSF Chimera, developed by the Resource for Biocomputing, Visualization, and Informatics at the University of California, San Francisco, with support from NIH P41-GM103311.

## SUPPLEMENTARY INFORMATION

**Supplemental Figure 1:**
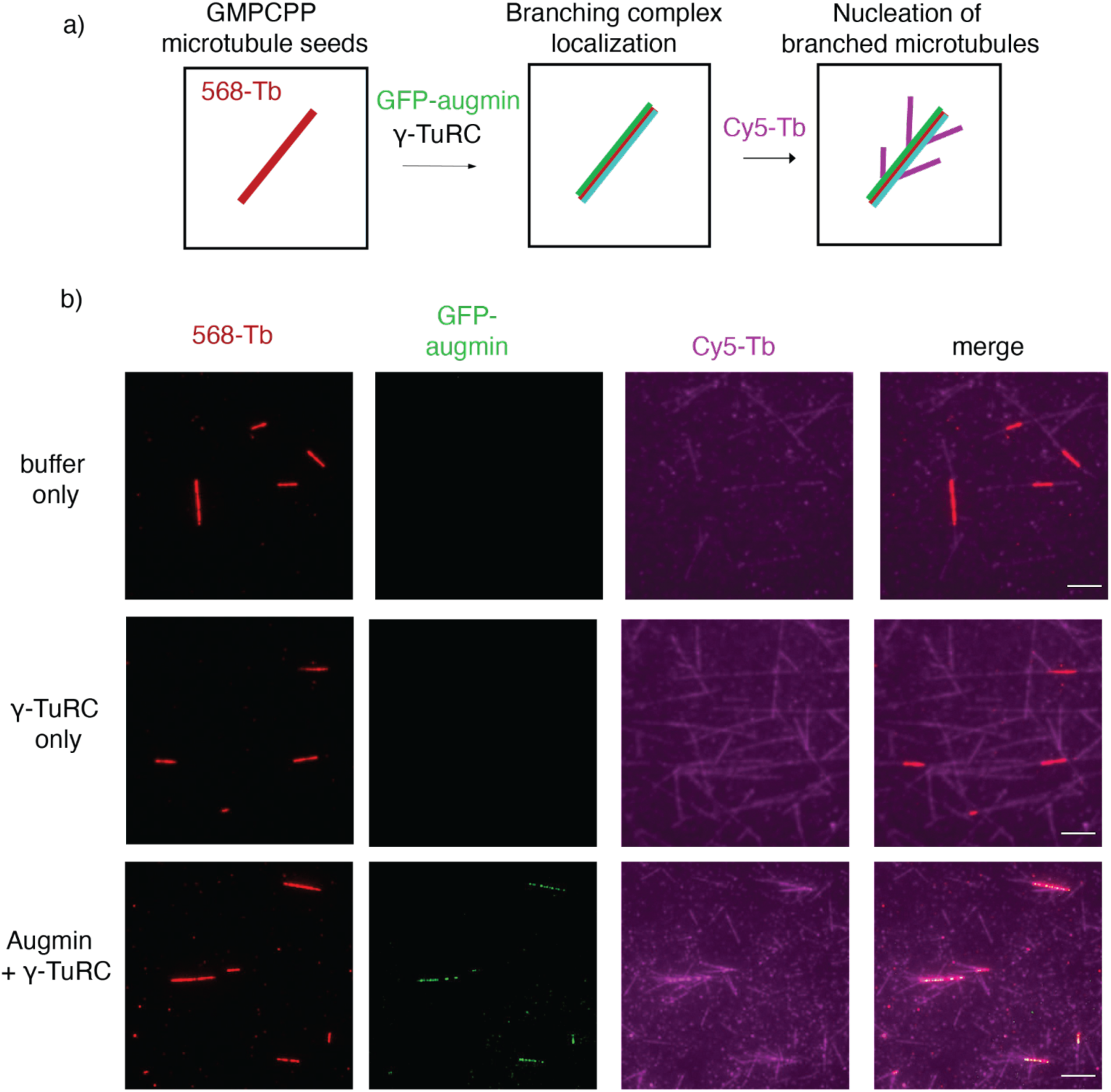
In vitro reconstitution of branching MT nucleation via TIRF microscopy. a. Schematic of in vitro branching reconstitution. Single cycled, GMPCPP stabilized MT seeds are attached to biotin-PEG functionalized glass cover slips via neutravidin and incorporation of biotin into the MT seeds. Following this, the branching factor complex containing augmin and γ- TuRC are bound to MT seeds. Finally, Cy5 labeled soluble tubulin was flowed into the reaction chambers to nucleate new MTs. b. Representative TIRF microscopy images from in vitro branching reconstitution performed with buffer only (top row), γ-TuRC only (middle row), and augmin/γ-TuRC (bottom row). MT seeds are visualized with Alexa-568 (red, first column), augmin is visualized by GFP labeling (green, 2nd column), and soluble tubulin is visualized by Cy5 labeling (3rd column). Merged images are shown in the last column. Only branching machinery containing augmin and γ-TuRC can effectively recruit soluble tubulin to the MT and initiate new branched MTs. Scale bars indicate 5 μM.

**Supplemental Figure 2:**
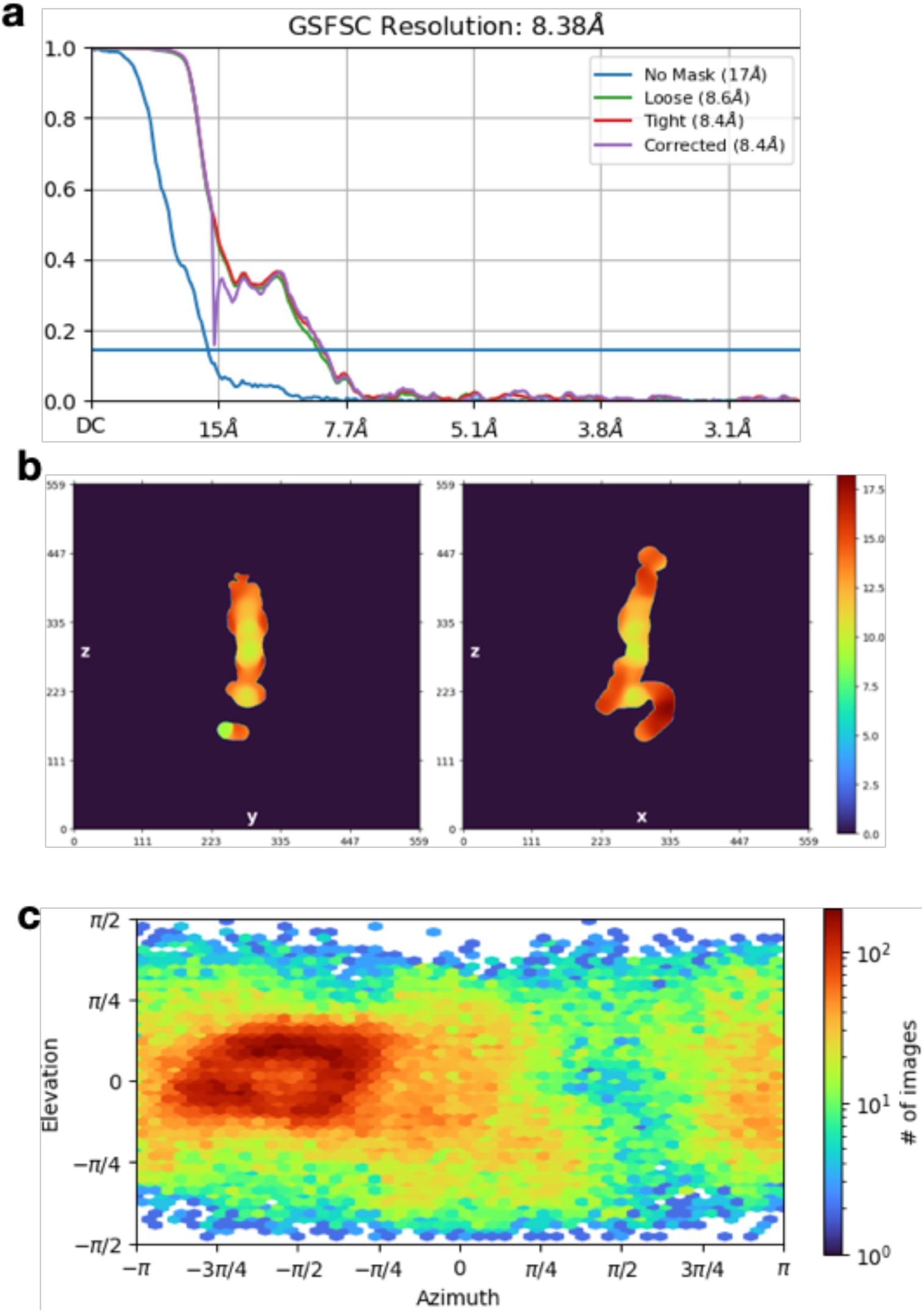
Cryo-EM map quality and features. a. Gold-standard FSC curves for determining cryo-EM map resolution are displayed, showing a resolution cut-off of 8.4 Å b. Local resolution determination of the cryo-EM map, demonstrating highest resolution in the center of the T-III stalk, and the lowest resolutions at the tips of T-III and T-II. c. Viewing direction distribution of particles in final map, showing a preferred orientation cluster at -π/2.

**Supplemental Figure 3:**
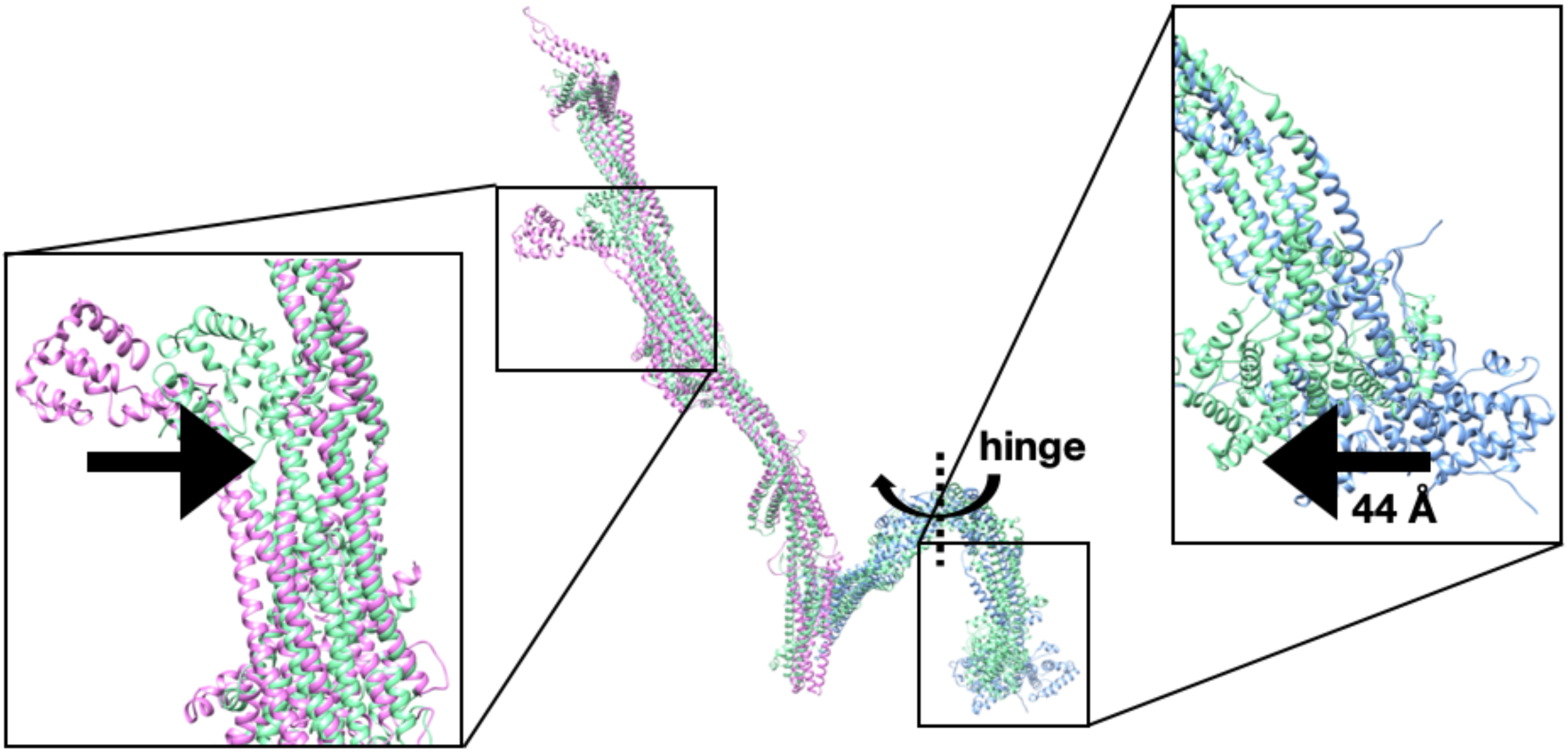
Fitting augmin model into cryo-EM map using MDFF. Molecular dynamic flexible fitting (MDFF) of the AlphaFold-Multimer T-III (pink) and T-II (blue) models into the cryo-EM map resulted in a density-guided model (green). Large scale collective motions occurred in the extreme N-termini of Haus3 and Haus5 (inset at left), which moved closer to the T-III stalk. In addition, the N-terminus of T-II rotated substantially at the indicted hinge (inset at right), both closing the T-II arch somewhat and substantially rotating the Haus6 and Haus7 calponin homology domains.

**Supplemental Figure 4:**
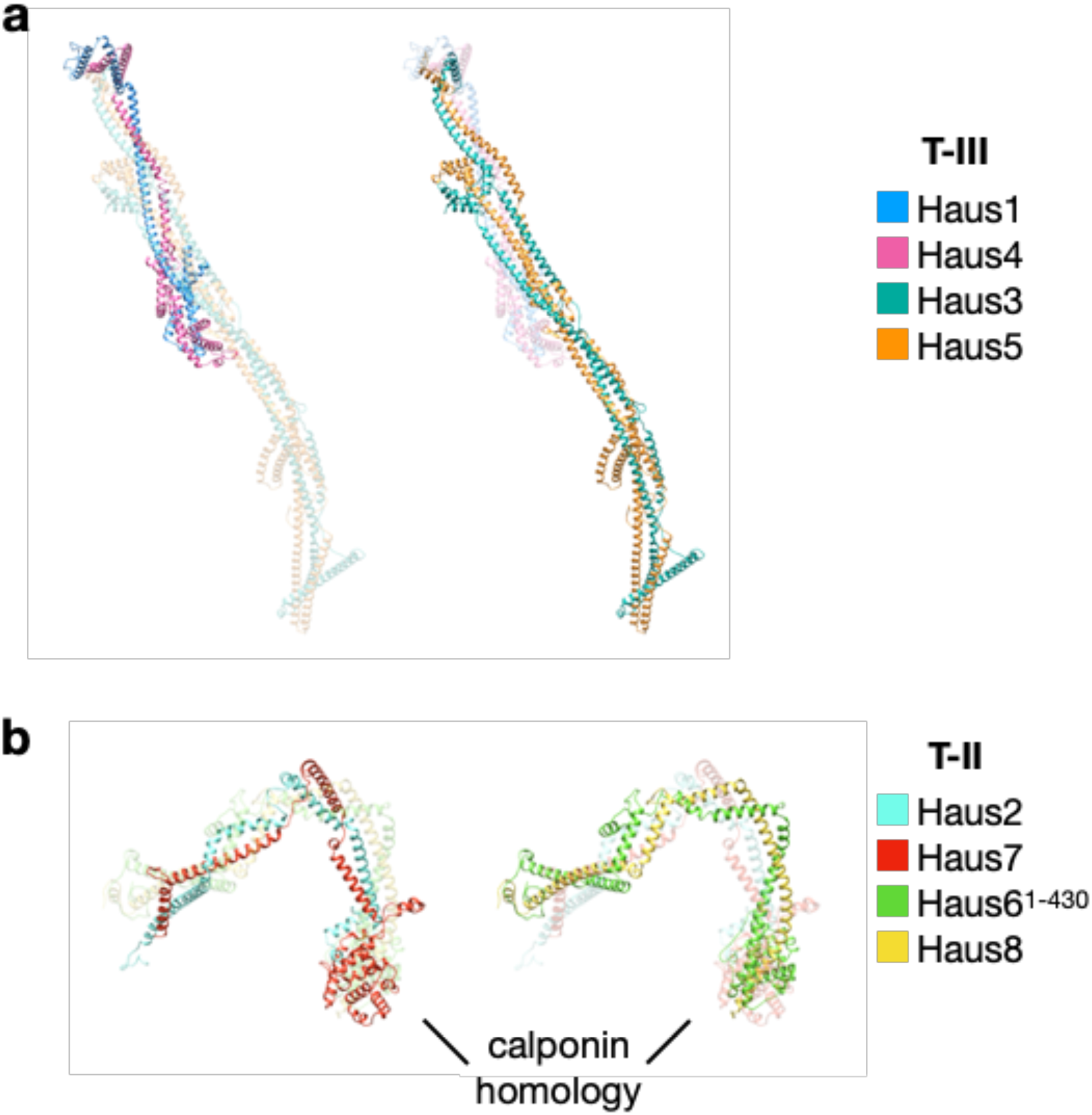
Coiled-coil architecture of augmin tetramers. a. T-III is a dimer comprised of two dimers: highlighted at left, the paralogs Haus1 (blue) and Haus4 (pink); highlighted at right, the much larger paralogs Haus3 (green) and Haus5 (orange). b. T-II is a dimer comprised of two dimers: highlighted at left, Haus6 (green) and Haus8 (yellow); highlighted at right, Haus2 (cyan) and Haus7 (red). Haus6 and Haus7 both contain calponin homology domains at their N-termini, marking the two as paralogs of one another.

**Supplemental Figure 5:**
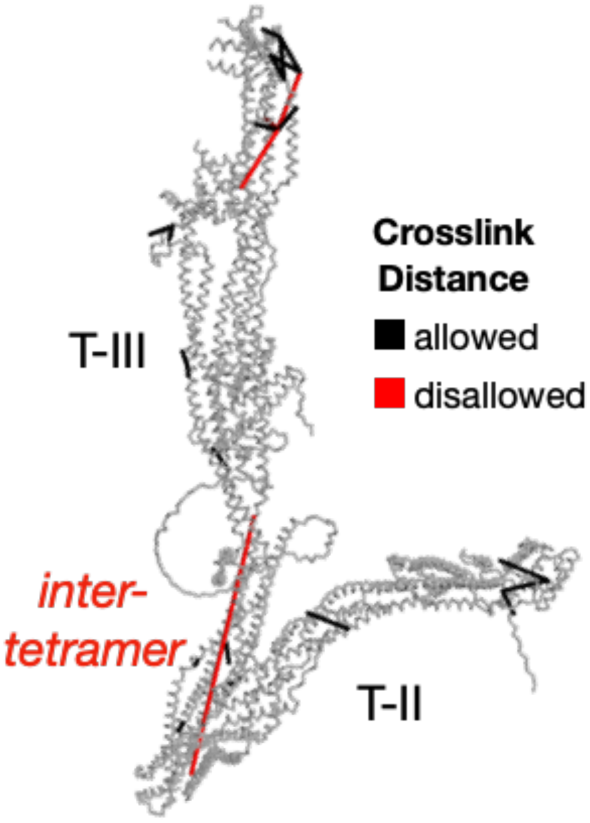
Model validation of *D. melanogaster* augmin using cross-linking mass spectrometry. Twenty-six high-confidence BS3 inter-subunit primary amine crosslinks that had previously been experimentally determined for *D. melanogaster* augmin were mapped to the integrated *D. melanogaster* augmin complex model. Inter-nitrogen distances within the allowed spacer arm length of 24 Å are indicated in black, whereas disallowed crosslinks of more than 24 Å are indicated in red. 22 out of 26 crosslinks are allowed. Crosslinking mass spectrometry data was derived from ^29^. Only crosslinks observed more than once are displayed.

**Supplemental Figure 6:**
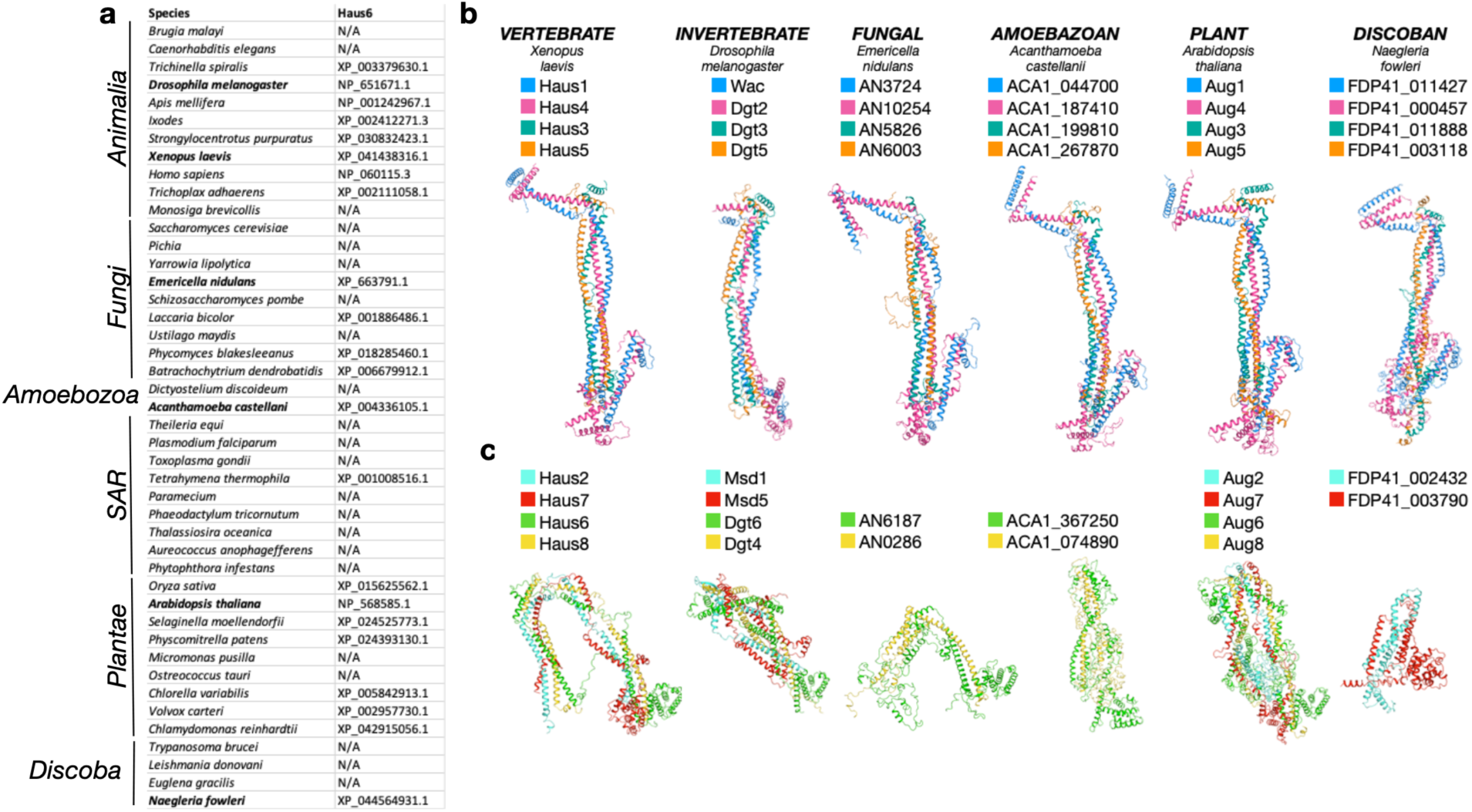
Conservation of the augmin complex across the eukaryotic kingdom. **a.** Results of sequence-based search for Haus6 orthologs across eukaryotes, with species listed on left and ortholog UniProt ID on right (species where no ortholog was detected are marked with N/A). Species selected for further modeling are marked in boldface. **b.** Augmin subunits in six representative eukaryotic species were identified either by prior independent work—*E. nidulans*^31^ and *A. thaliana*^16^—or by bioinformatic search of the assembled predicted proteome. Complete T-III^core^ complexes and partial or complete T-II complexes were modeled by AlphaFold-Multimer.

